# Temperature and Light Effects on Germination of *Peronospora effusa* Sporangia

**DOI:** 10.1101/163923

**Authors:** Robin Alan Choudhury, Neil McRoberts

## Abstract

Spinach downy mildew, caused by the biotrophic oomycete *Peronospora effusa*, is an economically important disease that is found in all spinach growing regions of the US. To effectively predict disease risk we need to understand the response of *P. effusa* to different environmental conditions. We conducted several germination assays, exposing *P. effusa* sporangia to different temperature and lighting conditions. Between 5 and 25°C under constant darkness, germination of *P. effusa* sporangia on water agar declined. These results were qualitatively different from a previous study of *P. effusa* germination that found a bimodal response curve, with increased germination at lower and higher temperatures. Time course studies revealed that sporangia consistently germinated within the first twelve hours of plating, regardless of incubation temperature. Sporangia exposed to blue light had significantly reduced germination when compared with those exposed to red, yellow, or no light. Light intensity and color significantly impacted germination, although the effect of color varied by light intensity.

---

Spinach downy mildew, caused by the biotrophic oomycete *Peronospora effusa* (Grev.) Rabenh (formerly *P. farinosa* f. sp. *spinaciae*) (Choi et al. 2007), is the most important threat to spinach production worldwide (Correll et al. 2011). Spinach downy mildew, like many other downy mildew diseases, typically occurs in cool, moist environments (Choudhury et al. 2016a). Coastal California frequently has cool morning temperatures and thick layers of fog and low clouds (Choudhury et al. 2016a) and produces approximately 40% of the annual US fresh market crop of spinach (USDA NASS 2016). While these cool temperatures can persist well into the afternoon for some portions of the region, more southern areas of coastal California frequently have warm, dry afternoon conditions for much of the year. Outbreaks of spinach downy mildew frequently occur despite these warmer and drier conditions, and can lead to large crop failures (Choudhury et al. 2016a). These disease outbreaks in warmer regions have many growers concerned that *P. effusa* may display phenotypic plasticity, and have a wider range of optimal environmental conditions for growth compared with other downy mildew pathogens.

To effectively predict disease risk we need to understand the response of *P. effusa* to different environmental conditions. However, compared with many other downy mildew pathogens, relatively little research has been done on the optimal environmental conditions for growth and sporulation of *P.effusa*. Recent field studies have revealed that environmental variables play a large role in the epidemiology of this disease (Choudhury et al. 2016a; Choudhury et al. 2016b). However, without understanding how individual environmental factors impact different stages of the pathogen life cycle, it is difficult to create an accurate risk prediction system. These controlled environment studies help to clarify how environmental variables affect the pathogens growth, reproduction and spread in isolation, without other confounding environmental effects.

An early report of germination rates of a *P. effusa* isolate from California suggested that the overall temperature range for germination was roughly the same, but that the optimal temperatures for germination were between 10 and 15°C (Cook 1937). While this report provided a glimpse into the optimal conditions for *P. effusa*, it did not provide a graph or table as to specific response to temperature. A later study from the Netherlands suggested that spinach downy mildew isolates had a bimodal reaction to increasing temperatures (Frinking et al. 1981). They found that *P. effusa* sporangia germinated between the temperatures 0 and 30°C. Interestingly, they found sporangial germination peaked at both 10 and 25°C, with a reduction of germination around 20°C. A bimodal temperature response in germination could help to explain how disease outbreaks can occur in fairly warm and dry regions as well as cooler regions. With a bimodal germination pattern, the pathogen could locally adapt to the local environmental conditions.

While the data for how *P. effusa* is affected by sub-optimal environmental conditions may be sparse, there is a wealth of knowledge regarding how other closely related species are affected by increasing temperatures. (Leach 1931) found that the beet downy mildew pathogen *P. schachtii* could germinate between 0 and 30°C, but had an optimal germination between 4 and 10°C. Interestingly, this optimal temperature range was shifted to 2 to 8°C when the isolate was grown in greenhouse conditions. Cook (1932) found that the onion downy mildew pathogen *P. destructor* had an optimal germination rate between 3 and 14°C, but that germination steeply declined in warmer temperatures. The cabbage downy mildew pathogen *Hyaloperonospora parasitica* seemed to have an optimal germination rate between 15 and 25°C, although it could germinate anywhere between 10 and 35°C (Achar 1998).

While understanding the overall range of optimal temperatures is important for future disease prediction efforts, understanding the timescale of sporangial germination is critical to the timing of protective fungicide sprays (Caffi et al. 2010). Most fungicides are protective, and help to inhibit the growth of the pathogen before it has entered the plant. Correct timing of fungicide applications can help to prevent the establishment of fungal pathogens in host plants. However, once a pathogen has entered the plant and created and infection center, protective fungicides typically have no useful effects. Timing the fungicide application to ensure that the susceptible germination tube is exposed is critical to prevent infection and crop failure.

Some fungi and oomycetes experience both fungicidal and fungistatic effects of sub-optimal environmental conditions. The onion downy mildew pathogen *P. destructor* had a delayed and reduced germination effect when germinated at increasing temperatures (Cook 1932). However, while the beet downy mildew pathogen *P. schachtii* had reduced germination at increased temperatures, most germination had occurred within the first 4 hours after removal from the leaf (Leach 1931). Similarly, while cabbage downy mildew pathogen *Hyaloperonospora parasitica* did experience reduced germination outside of its optimal growth temperatures, there was no noticeable delay to germination (Achar 1998). Understanding how sup-optimal temperature delays germination can help determine when the pathogen will be most susceptible to protective fungicides.

Quality and quantity of light can have dramatic effects on many fungal and oomycete organisms. The overall quantity and period of light can drive different processes of oomycete biology, including sporulation and infection (Cohen et al. 2013). In addition to the effects from the total amount of light exposure, many oomycetes are affected by specific spectra of light. Some oomycetes are inhibited by blue light, which can reduce sporulation and growth (Cohen et al. 1975; Cohen 1976; Cohen and Eyal 1977). Interestingly, a recent report suggests that the closely related pathogen *P. belbahrii* is most negatively affected by red light, which strongly suppresses sporulation (Y. Cohen et al. 2013). Some basil growers have taken advantage of this detrimental effect on sporulation by using red light in susceptible basil greenhouses, thereby reducing downy mildew epidemic severity. Understanding how *P.effusa* is affected by different qualities and quantities of light may help to guide future control strategies for spinach downy mildew.

In this study, we exposed *P. effusa* sporangia to different temperature and lighting conditions and monitored for detrimental effects on germination. The goals of this study were to understand: (1) how California isolates of *P. effusa* are affected by temperature, (2) how temperature affects the timescale of sporangial germination, and (3) how different qualities and quantities of light affect sporangial germination.

## MATERIALS AND METHODS

### *Peronospora effusa* source

Six isolates of *P. effusa* were collected from diseased fields in the Salinas Valley in 2015 and 2016 (Table 1). Isolates were considered distinct if they were localized in small disease outbreaks (<20cm) that were spatially separated from other outbreaks, therefore likely arising from the expansion of a single infection event. Approximately 20 diseased leaves from each isolate were collected into sterile bag containers, and two bags were collected from each outbreak source. Bags were sealed and stored at 4°C in a cooler until processing. Sporangia were removed from infected leaves by applying 50ml of chilled sterile distilled water into the sealed bags and shaking vigorously for five minutes. The resulting spore suspension was standardized to 1×10^5^ spores/ml using a haemocytometer. Spore suspensions were maintained on ice while being prepared for experimental use.

**TABLE 1:** Information on the source of *Peronospora effusa* isolates used in this study.

| Isolate # | Variety | Location | Collection Month | Temperature<sup>a</sup> |
| --- | --- | --- | --- | --- |
| DNM001 | Violin | King City, CA | June, 2015 | 18.1 |
| DNM002 | Platypus | Hollister, CA | July, 2015 | 19.5 |
| DNM003 | Carmel | Hollister, CA | July, 2015 | 19.5 |
| DNM004 | Emilia | King City, CA | May, 2016 | 15.5 |
| DNM005 | Emilia | King City, CA | May, 2016 | 15.5 |
| DNM006 | Viroflay | Watsonville, CA | July, 2016 | 13.7 |

<sup>a</sup> Average temperature during the month of collection of isolate source location

### Effect of temperature on *P. effusa* germination

150ul of *P. effusa* 10^5^ sporangia/ml spore suspension were plated and spread uniformly on2% water agar on 100×15mm plates. Plates were immediately stored in crisper containers, and the containers were placed into incubation chambers maintained at 5, 10, 15, 20, and 25°C in darkness. Incubation chamber temperatures were confirmed using analog thermometers. Ten plates were used for each temperature treatment. At 24h after plating, crisper boxes were removed from incubation chambers. Plates were examined under a 10× light microscope for sporangial germination. One hundred sporangia from each plate were rated for germination status. Sporangia that had visibly developed sub-terminal or lateral germ tubes were considered germinated. Experiments were repeated twice.

The effect of temperature on sporangial germination over time was measured for *P. effusa* isolate DNM003. Plates were treated as described above, and sporangial germination was measured at 4, 8, 12, and 24 hours after plating. The experiment was repeated twice.

### Effect of light quality on *P. effusa* germination

Crisper boxes were fitted with red, yellow, green, and blue color correction gels (Cowboy Studio LLC., Texas, USA) to allow exposure to specific qualities of light. The bottoms and sides of the crispers were covered in aluminum foil to prevent ambient light. The crisper box used to test the effect of full dark on germination was completely covered with aluminum foil, and the crisper box used to test no filter was left unaltered.

**Supplementary Figure 1:**
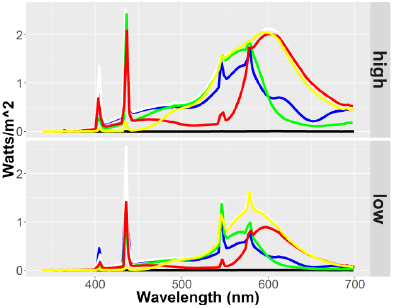
Quality and quantity of light exposed to *Peronospora effusa* isolate DNM003 sporangia. The unfiltered light source is indicated by the white line.

Seventy-five microliters of *P. effusa* isolate DNM003 spore suspension (10^5^ sporangia/ml) were plated and spread uniformly on 2% water agar on 60×15mm plates. Plates were laid in a single layer on the bottom of the crisper boxes, and boxes were immediately moved into a growth chamber (model E7, Conviron Ltd., Winnapeg, Canada). The effects of light intensity and temperature were tested alongside light quality in a fully factorial design. The growth chamber was set to either constant full fluorescent and incandescent lamps or constant half-power fluorescent lamps, at either 4° or 18° C. The plates were removed and measured for sporangial germination as described above at 24h after plating. All experiments were repeated twice. The quality and quantity of light was measured within the sealed crisper boxes using a spectrometer (Black Comet model, StellarNet, Florida, USA) using the SpectraWiz software (StellarNet, Florida, USA).

### Statistical analysis

Means comparisons between different treatment temperatures, lighting conditions, and time periods were completed using Fisher’s least significant difference test, implemented using the R v3.2.4 statistical programming language (R Core Team 2016) and the LSD.test function in the package agricolae v1.2-3 (Mendiburu 2015). A linear regression analysis of the sporangial germination percentage against temperature was used to estimate the effect of temperature on sporangial germination for each individual isolate.

## RESULTS

### Effect of temperature on *P. effusa* germination

Across all isolates, there was a consistent decline in germination rate as the exposure temperature increased (Fig. 1).

**Fig. 1:**
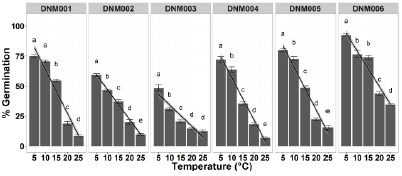
Mean and standard error of percentage germination of different *Peronospora effusa* isolates over different temperatures. Different letters represent significant differences between treatment temperatures (Fisher’s least significant difference test, α = 0.05).

Results of the Fisher’s least significant difference test suggest that while all tested *P. effusa* isolates had declining rates of germination as temperature increased, the effects were not uniform. Isolate DNM001 had no significant difference in germination rate between 5 and 10°C, isolate DNM003 had no significant difference in germination rate between 20 and 25°C, and isolate DNM006 had no significant difference in germination rate between 10 and 15°C. None of the tested isolates had a bimodal temperature response as was previously reported in Frinking et al. (1981). Results of the linear regression analyses suggest that sporangial germination declines by approximately 1.8 to 3.7% for every additional unit increase in temperature (Table 2). This relationship was strong and consistent across isolates.

**TABLE 2:** summary coefficients from the linear regression analyses of percentage germination of *P. effusa* sporangia fitted to temperature effects by isolate.

| Isolate | Intercept | Slope | R<sup>2</sup> Coefficient | p value |
| --- | --- | --- | --- | --- |
| DNM001 | 100.59 | -3.67 | 0.899 | < 2.2e-16 |
| DNM002 | 72.03 | -2.50 | 0.879 | < 2.2e-16 |
| DNM003 | 52.02 | -1.78 | 0.711 | < 2.2e-16 |
| DNM004 | 91.95 | -3.51 | 0.895 | < 2.2e-16 |
| DNM005 | 101.62 | -3.59 | 0.907 | < 2.2e-16 |
| DNM006 | 108.77 | -2.97 | 0.835 | < 2.2e-16 |

*Peronospora effusa* sporangia typically germinated within the first 12 hours of plating. There were no significant increases in germination between 5 and 20°C after 12 hours after plating (Fig. 2). At 25°C, there was a significant increase in germination rate after 12 hours. At 5°C, there was no significant increase in germination rate after 8 hours.

**Fig. 2:**
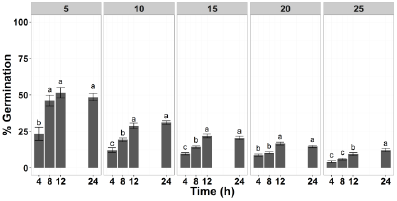
Mean and standard error of percentage germination of *Peronospora effusa* isolate DNM003 sporangia as measured at 4, 8, 12, and 24h after plating. Different letters represent significant differences between time points (Fisher’s least significant difference test, α = 0.05).

### Effect of light quality on *P. effusa* germination

Unfiltered light significantly reduced *P. effusa* germination compared with the no light control at full light intensity, but not at low light intensity (Fig. 3). Across both temperatures and light intensities, exposure to blue light consistently reduced germination when compared with the no light control. Green light significantly reduced germination compared with the no light control at full light intensity, but not at low light intensity. Yellow light significantly increased germination at full light intensity and low light intensity at 18°C compared with the no light control, but had no significant effect at low light intensity at 4°C. Red light had no significant effect on germination compared with the no light control. The wavelength spectra of the crisper boxes used with color correction gels shows non-specific wavelengths outside of the target spectra (Supplementary Fig. 1).

**Fig. 3:**
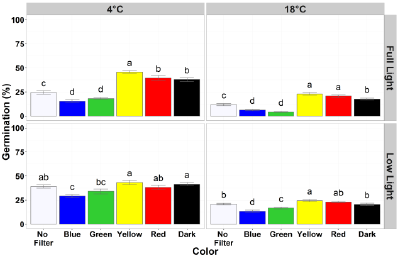
Mean and standard error of percentage germination of *Peronospora effusa* isolate DNM003 sporangia exposed to different temperature and lighting conditions. Different letters represent significant differences between treatment colors (Fisher’s least significant difference test, α = 0.05).

## DISCUSSION

In this study, *P. effusa* sporangia were exposed to different environmental conditions to observe their effects on germination. While there were some isolate specific effects of temperature, all isolates had reduced germination when exposed to increasing temperatures. A linear regression analysis suggests that for every unit increase in temperature, *P. effusa* germination declines by approximately 1.8-3.7%. While temperature reduced overall germination of sporangia, it did not have a consistent significant detrimental effect on the time until germination. Blue light significantly reduced germination of sporangia at all temperature and lighting conditions, yellow light significantly increased sporangial germination in full light, and red light had no significant impact on sporangial germination.

A previous report from the Netherlands found a bimodal response of *P. effusa* sporangia to increasing temperature. We did not observe a similar temperature response in the six California isolates that we characterized. Based on these results, it is unlikely that environmental adaption helps to explain the epidemics observed in the warmer regions of coastal California (Choudhury et al. 2016). It is possible that those epidemics are driven by the increased availability of hosts rather than conducive environmental conditions. It is also possible that the conducive conditions that occur in the early morning before the increase of heat allow for pathogen infection. The results of our study more closely align with those of an earlier report (Cook 1932) that suggested that the optimal temperature for *P. effusa* sporangial germination was between 10 and 15°C.

In other similar controlled environment studies, isolates of the same species can sometimes have different responses to similar environmental conditions. A study comparing the effects of temperature on germination, infection, and sporulation of two geographically distinct isolates of the maize downy mildew pathogen *Peronosclerospora sorghi* found that there were significant differences between isolates from Asia and North America (Bonde et al. 1985). A similar study of the temperature response of two rose-infected downy mildew species, *Peronospora sparsa* and *P. rubra*, found that some isolates had increased germination at higher temperatures (Breese et al. 1994). This phenotypic plasticity between isolates may help with adaptation to new environments. It is possible that the population of *P. effusa* isolates characterized in the previous study had an inherently different response to environmental conditions, and were able to withstand exposure to a wider temperature range more readily than our California isolates. Further study of a broader population of *P. effusa* isolates would help to characterize any regional adaption response to environmental conditions.

This study found that there was a fairly consistent time until germination for *P. effusa* sporangia. A study of the effect of temperature on germination in the closely related beet pathogen *P. schachtii* found that while temperature reduced germination, there was no delay in the timing of germination (Leach 1931). Another study on the onion downy mildew pathogen *P. destructor* found that increased temperature delayed sporangial germination (Cook 1932). It is possible that the time until germ tube development and the fungistatic effects of temperature may be genetically determined, with more closely related species having a similar response.

A previous study used empirical weather and spore trapping data to predict environmental effects on *P. effusa* spore capture (Choudhury et al. 2016). This current study found that increased temperatures and reduced winds were correlated with increased spore copy number detection using rotating arm spore traps. While these results may seem conflicting, it is likely that different environmental conditions guide germination and sporulation. This distinction between the conditions necessary for germination, infection and sporulation has been seen before in other downy mildew pathogens. Arauz et al. (2010) showed in controlled environment studies that there was an approximate 5°C gap in the optimal temperatures necessary for germination of *Pseudoperonospora cubensis*, the cucurbit downy mildew pathogen. Future controlled environment studies focused on the conditions required for *Peronospora effusa* infection and sporulation are necessary to predict the entire pathogen life cycle.

Previous studies have found that light quality and quantity can have different effects on downy mildew pathogen biology. We found that blue light reduced germination of sporangia, while yellow light promoted germination. Cohen et al. (1975) had similar results, with an inhibitory effect of blue light on production of sporangia for the potato late blight pathogen *Phytophthora infestans*. While they did not investigate the effects of yellow light, they also found that red light had no inhibitory effects on sporulation. A similar pair of studies found that blue light inhibited sporulation in *Peronospora tabacina* and *Pseudoperonospora cubensis*, the tobacco and cucurbit downy mildew pathogens, respectively (Cohen 1976; Cohen and Eyal 1977).

These effects of light on pathogen growth might tie in with inherent circadian rhythms within *Peronospora effusa*. A recent study found that the *Arabadopsis thaliana* downy mildew pathogen *Hyaloperonospora arabadopsidis* had significantly more successful infection events during evening hours rather than daylight hours (Wang et al. 2011). Although we tested our germination assays using water agar, these patterns may inherently persist even in the absence of a host. This effect could lead to experimental variation even within a single isolate.

This study highlighted the potential differences in the environmental response of different isolates of the same species. Our findings show that the germination rates of Californian isolates of *Peronospora effusa* were typically negatively affected by increasing temperatures. While this study helps to lay the groundwork for a risk prediction model, germination is only a portion of the *P. effusa* lifecycle. Further controlled environment studies investigating growth, sporulation, and dispersal are needed to fully predict this disease.

## ACKNOWLEDGEMENTS

We thank the California Leafy Greens Research Program (CLGRP) for funding this research. We thank Mr. Steven T. Koike for help in locating isolates and technical recommendations and Drs. James Correll, R. Michael Davis, Sierra Hartney and Johan Leveau for helpful discussion. We thank Drs. Miki Okada and Julin Maloof for training and use of spectrophotometer.

## LITERATURE CITED

Achar, P. (1998). Effects of temperature on germination of Peronospora parasitica conidia and infection of Brassica oleracea. Journal of Phytopathology, 146(2-3), 137-141.

Arauz, L., Neufeld, K., Lloyd, A., & Ojiambo, P. (2010). Quantitative models for germination and infection of Pseudoperonospora cubensis in response to temperature and duration of leaf wetness. Phytopathology, 100(9), 959-967.

Bonde, M., Peterson, G., & Duck, N. (1985). Effects of temperature on sporulation, conidial germination, and infection of maize by Peronosclerospora sorghi from different geographical areas. Phytopathology, 75(1), 122-126.

Breese, W. A., Shattock, R., Williamson, B., & Hackett, C. (1994). In vitro spore germination and infection of cultivars of Rubus and Rosa by downy mildews from both hosts. Annals of Applied Biology, 125(1), 73-85.

Caffi, T., Rossi, V., & Bugiani, R. (2010). Evaluation of a warning system for controlling primary infections of grapevine downy mildew. Plant Disease, 94(6), 709-716.

Choi, Y. J., Hong, S. B., & Shin, H. D. (2007). Re-consideration of Peronospora farinosa infecting Spinacia oleracea as distinct species, Peronospora effusa. Mycol Res, 111(Pt 4), 381-391, doi:10.1016/j.mycres.2007.02.003.

Choudhury, R., Koike, S., Fox, A., Anchieta, A., Subbarao, K., Klosterman, S., et al. (2016). Season-Long Dynamics of Spinach Downy Mildew Determined by Spore Trapping and Disease Incidence. Phytopathology, 106(11), 1311-1318.

Cohen (1976). Interacting effects of light and temperature on sporulation of Peronospora tabacina on tobacco leaves. Australian Journal of Biological Sciences, 29(3), 281-290.

Cohen, & Eyal, H. (1977). Growth and differentiation of sporangia and sporangiophores of Psudoperonospora cubensis on cucumber cotyledons under various combinations of light and temperature. Physiological Plant Pathology, 10(2), 93IN197-196IN2103.

Cohen, Eyal, H., & Sadon, T. (1975). Light-induced inhibition of sporangial formation of Phytophthora infestans on potato leaves. Canadian Journal of Botany, 53(22), 2680-2686.

Cohen, Y., Vaknin, M., Ben-Naim, Y., & Rubin, A. E. (2013). Light suppresses sporulation and epidemics of Peronospora belbahrii. PLoS One, 8(11), e81282, doi:10.1371/journal.pone.0081282.

Cook, H. T. (1932). Studies On The Downy Mildew Of Onions, And The Casual Organism, Peronospora Destructor (Berk.) Caspary. Cornell University Agricultural Experiment Station memoir, 143.

Cook, H. T. (1937). Germination of conidia of Peronospora effusa from spinach. Phytopathology, 27, 124.

Correll, J., Bluhm, B., Feng, C., Lamour, K., Du Toit, L., & Koike, S. (2011). Spinach: better management of downy mildew and white rust through genomics. European Journal of Plant Pathology, 129(2), 193-205.

Frinking, H., Geerds, C., & Meerman, F. (1981). Germination of Peronospora farinosa f. sp. spinaciae conidia: a two-topped temperature curve. European Journal of Plant Pathology, 87(4), 163-165.

Leach, L. D. (1931). Downy mildew of the Beet, caused by Perono-spora schachtii Fuckel. Hilgardia, 6(7), 203-251.

Wang, W., Barnaby, J. Y., Tada, Y., Li, H., Tör, M., Caldelari, D., et al. (2011). Timing of plant immune responses by a central circadian regulator. Nature, 470(7332), 110-114.

